# Accelerating Identification of Chromatin Accessibility from noisy ATAC-seq Data using Modern CPUs

**DOI:** 10.1101/2021.09.28.462099

**Authors:** Narendra Chaudhary, Sanchit Misra, Dhiraj Kalamkar, Alexander Heinecke, Evangelos Georganas, Barukh Ziv, Menachem Adelman, Bharat Kaul

**Affiliations:** Intel Labs, Bangalore, India; Intel Labs, Santa Clara, California, USA; Intel Corporation, Haifa, Israel

**Keywords:** chromatin accessibility, ATAC-seq, deep learning, 1D convolutions, architecture-aware optimizations, SIMD, BF16

## Abstract

Identifying accessible chromatin regions is a fundamental problem in epigenomics with ATAC-seq being a commonly used assay. Exponential rise in single cell ATAC-seq experiments has made it critical to accelerate processing of ATAC-seq data. ATAC-seq data can have a low signal-to-noise ratio for various reasons including low coverage or low cell count. To denoise and identify accessible chromatin regions from noisy ATAC-seq data, use of deep learning on 1D data – using large filter sizes, long tensor widths, and/or dilation - has recently been proposed. Here, we present ways to accelerate the end-to-end training performance of these deep learning based methods using CPUs. We evaluate our approach on the recently released AtacWorks toolkit. Compared to an Nvidia DGX-1 box with 8 V100 GPUs, we get up to 2.27× speedup using just 16 CPU sockets. To achieve this, we build an efficient 1D dilated convolution layer and demonstrate reduced precision (BFloat16) training.

## Background

Human DNA has approximately three billion base pairs, but most of it is inaccessible due to its organization in the chromatin material. Chromatin material consists of folded-up DNA and proteins. Depending upon the molecular folding, different parts of DNA are accessible or inaccessible. Determining the accessibility level of different parts of the genome is important to understand their role in gene regulation. ATAC-seq is a commonly used technology that measures the accessibility level for each position in the DNA sequence. The ATAC-seq based method uses a Tn5 transposase molecule to tag segments of DNA that have higher accessibility. It uses next-generation sequencers to produce multiple sequencing reads from those segments. A higher number of reads from a particular region of the genome signifies higher accessibility. Therefore, the reads are mapped to the reference genome to create a one-dimensional (1D) array containing the number of reads mapped to each position in the reference genome. Subsequently, peak calling is performed on the 1D array to output accessible chromatin regions. In this paper, we focus on the peak calling problem.

ATAC-seq data can have a high signal-to-noise ratio for various reasons, including low coverage, low cell count, and low-quality data – making it difficult to perform accurate peak calling. It is especially true for single-cell ATAC-seq experiments. Given the exponentially increasing throughput and reducing cost of next-generation sequencing and the corresponding rise in ATAC-seq experiments, it is critical to develop accurate and high throughput methods for denoising ATAC-seq data and peak calling. To this end, several deep learning based methods have been proposed to perform denoising and peak detection from 1D sequencing data [1–8]. These deep learning based methods typically use deep convolutional neural networks (CNNs). CNNs have been widely used in genomics applications, such as, variant calling [9], gene expression prediction [10], and inferring gene relationships from single-cell expression data [11]. In CNN models, input data passes through a series of convolution layers.

For denoising ATAC-Seq data and peak calling in particular, two recent approaches are especially noteworthy. Rai et al., used a CNN architecture [12] with 1D convolution layers to upscale ATAC-seq [13] data. The AtacWorks toolkit from Lal et al. [2] that is based on 1D convolution layers was used for simultaneous denoising and peak calling at the base-pair level resolution from low-coverage, low cell count, or low-quality ATAC-seq data. AtacWorks was also able to denoise single-cell ATAC-seq data. The AtacWorks model uses a ResNet CNN architecture with multiple 1D convolution layers with dilated convolutions [14], large filter sizes, and long tensor widths. Deep CNNs equipped with 1D dilated convolution layers have long receptive field sizes, especially if they contain large filters. Genome sequencing data can have relationships spanning long distances along the genome, and longer receptive fields can capture this information. Hence, we expect that deep CNNs, similar to AtacWorks, will continue to find applications in research problems that use genomics sequencing data. The primary time-consuming module in the above two approaches is the convolution layer over 1D data. However, existing implementations of the 1D convolution layer for CPUs and GPUs fail to efficiently use the underlying hardware especially in the case of large filter sizes, long tensor widths, and dilation.

### Our contributions

In this work, we accelerate denoising and identifying chromatin accessibility by improving the training time of deep CNNs – that use 1D convolutions – by utilizing multi-core CPUs. We demonstrate the performance benefits of our approach on the AtacWorks toolkit. Prior to our work, the most efficient CNN implementation available for the CPUs was from oneDNN, an open-source performance library of key computational modules for deep learning applications. However, oneDNN failed to efficiently utilize the underlying hardware for 1D convolutions with dilation and large filters that are present in the AtacWorks model. We employ the following methods to achieve acceleration by improving the computational efficiency.

1. We create an efficient and generic 1D dilated convolution layer [15] in the PyTorch framework. To implement our 1D convolution layer, we employ tensor processing primitives (TPP) [16] from the LIBXSMM library [17]. TPPs use jitting and efficient algorithms to provide extremely fast implementations of key deep learning primitives on CPUs. It has been shown that most of the popular deep learning algorithms can be implemented using just one primitive as the basic building block – batch reduce generalized matrix multiplication (BRGEMM). Our implementation utilizes the BRGEMM TPP [18] and cache blocking to build the most efficient implementation of 1D convolutions with dilations and large filter sizes on modern CPUs.
2. We employ optimizations for data loading, multithreading, and scaling for efficient training on single-socket and multiple sockets of CPUs.
3. Further, we demonstrate the first-ever end-to-end training of the AtacWorks CNN model using reduced precision of BF16 without any loss of accuracy. Prior to this work, the AtacWorks model was only shown to work with single-precision floating-point (FP32) numbers. For the BF16 based implementation, we leverage AVX-512 BF16 instructions of Intel^®^ Xeon^®^ Cooper Lake (CPX) CPUs and implement all the steps of the end-to-end training workflow using BF16. Specifically, we create BF16 enabled 1D convolution layer, rectified linear unit (ReLU) layer, and avoid unnecessary tensor conversions. The use of reduced precision results in significantly faster training.

We report our experimental results for end-to-end training of AtacWorks CNN on various modern CPUs that are detailed in Supplementary Table S1. On a single-socket Intel^®^ Xeon^®^ Cascade Lake (CLX) CPU, our work achieves nearly 6.86× speedup over the corresponding Intel^®^ oneDNN library [19] based implementation. We also achieve nearly linear scaling with our implementation on multiple sockets of Intel^®^ Xeon^®^ Cascade/Cooper/Ice Lake CPUs. We demonstrate that our execution on multiple CPU sockets is significantly faster than the published results for DGX-1 [20] box with 8 V100s [2] without any loss of accuracy. For a fair comparison with DGX-1 box, we use CPU systems with similar power envelop. Compared to AtacWorks running on DGX-1 box [2], using 16 CPU sockets, we achieve up to 2.27× and up to 1.83× speedup for training with BF16 precision and FP32 precision, respectively.

## Results

In this section, we present the results of our end-to-end training experiments. First, we present end-to-end AtacWorks training time on single-socket CPUs. Subsequently, we scale our experiments by increasing the number of CPU sockets, dataset size, and ATAC-seq signal track size. We show multi-socket CPU scaling results and compare them with multi-GPU results published in [2].

### ATAC-seq Data and AtacWorks

In our experiments, unless explicitly stated otherwise, we use the following datasets and setup, which are same as the ones used in the AtacWorks paper [2]. We use the dsc-ATAC-seq dataset. This dataset contains single-cell ATAC-seq data from different types of human blood cells. First, 2400 monocytes are selected to make a clean high-coverage signal. After that, 50 monocytes are randomly sampled from these 2400 monocytes to make a noisy ATAC-seq signal. During training, the AtacWorks model attempts to learn 1) a mapping from noisy 50 cell signal to 2400 cell clean signal to use for denoising the 50 cell signal and 2) a model to call peaks. From the ATAC-seq dataset, we use chromosome 20 for validation, hold out chromosome 10 and use all other autosomes for training. We cut the full ATAC-seq signal into signal track segments of width 50000 and pad it on both sides by 5000 points to make it of width 60000. This way, the entire training set contains 32000 1D ATAC-seq signal track segments, and the validation set contains 1280 signal track segments.

The AtacWorks neural network uses a ResNet [21] (residual neural network) architecture that consists of multiple residual blocks. Figure 1 shows one such residual block that consists of three 1D dilated convolution layers followed by ReLU nonlinearities. AtacWorks utilizes five residual blocks to denoise the ATAC-seq signal and two additional residual blocks to call peaks. Multiple loss functions are used to train the AtacWorks network. Mean squared error (MSE) is used as a loss function on the denoised signal, and binary cross-entropy is used as a loss function on the peak array. The AUROC metric is used for the evaluation of the model. A trained AtacWorks model takes noisy ATAC-seq signal track segment as input and produces a corresponding denoised signal track segment along with a binary array of called peaks.

**Figure 1.**
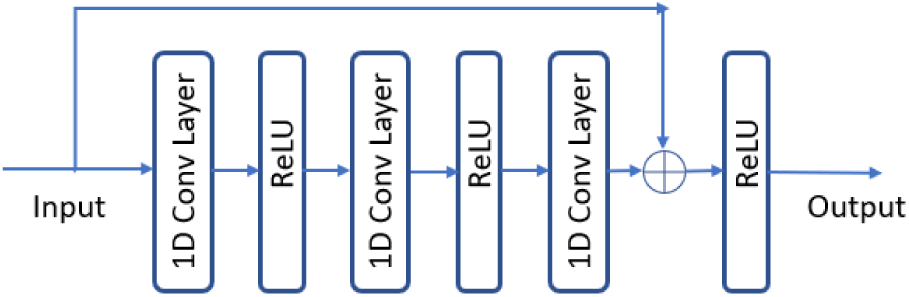
Residual block in AtacWorks CNN.

The core computational kernel of AtacWorks is a 1D dilated convolution layer. It takes more than 90% of the computation time during training. Figure 2 illustrates an example 1D dilated convolution layer along with parameters of filters, channels, input width, filter width, and dilation. Most convolution layers in AtacWorks use filter width of 51, dilation of 8, number of input channels as 15, and number of filters as 15. Generally, the input tensor width is 60000. Considering these parameters, we optimize our 1D convolution layer for large input widths, large filter widths, and dilation.

**Figure 2.**
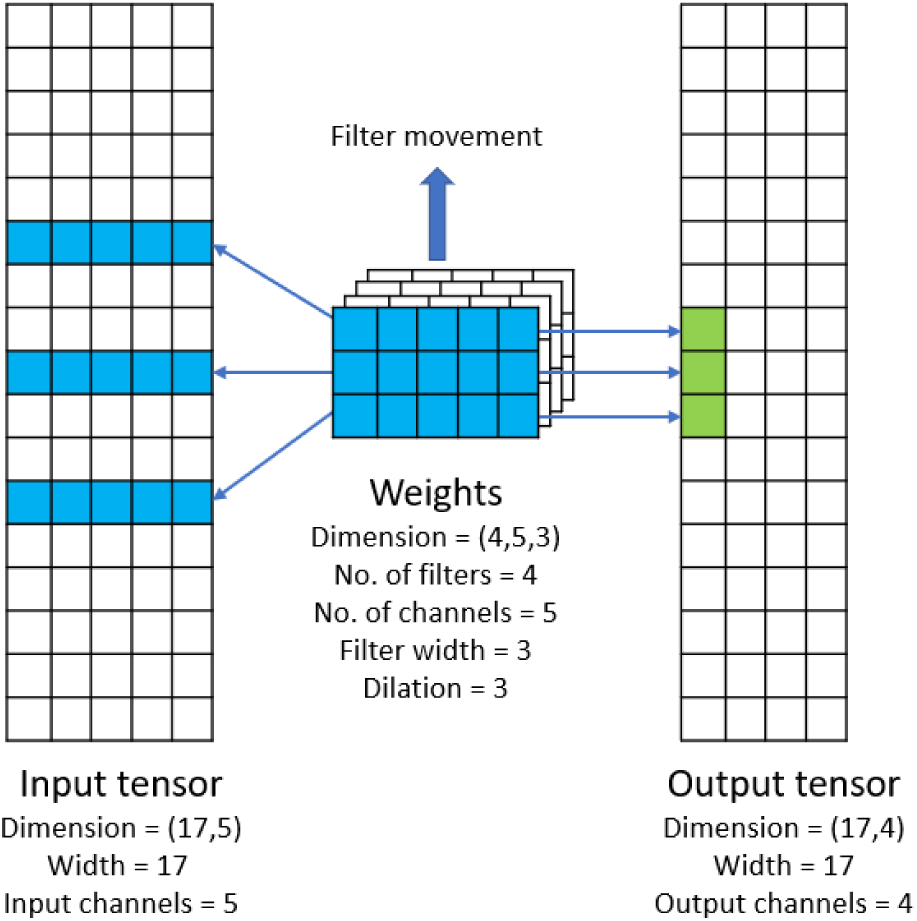
An example of 1D dilated convolution layer with parameters.

### System Details

In order to show the generality of our approach, we use four different CPUs for our experiments: i) Intel^®^ Xeon^®^ Platinum 8280 or Cascade Lake (CLX) with 28 cores per socket and AVX-512 FP32 support, ii) Intel^®^ Xeon^®^ Platinum 8380HL or Cooper Lake (CPX) with 28 cores per socket and AVX-512 FP32 and BF16 support, iii) Intel^®^ Xeon^®^ Platinum 8380 or Ice Lake (ICX) with 40 cores per socket and AVX-512 FP32 support, and iv) AMD EPYC 7742 (ROME) with 64-cores per socket and AVX2 FP32 support. We enable turbo for all cores during our experiments. More details of the systems used are provided in Supplementary Table S1.

### Our Implementation

The existing AtacWorks implementation uses the PyTorch framework, and its training source code only works with GPUs. We modify the source code of AtacWorks for training on single and multiple sockets of CPU. The PyTorch based implementation for 1D convolution has poor throughput. Therefore, we experiment with two separate implementations that replace the key computational modules of PyTorch framework’s backend with more performant alternatives. The first implementation uses optimized modules from the oneDNN library. On the other hand, the second one uses optimized modules that we implemented ourselves using LIBXSMM’s efficient tensor processing primitives and integrated into the PyTorch backend through Python extensions.

For single-socket training, our implementation uses one core on the socket for the data loader thread and the rest of the cores for neural network computation. Our multi-socket implementation uses the MPI communication protocol through the oneCCL library [22]. Therefore, for multi-socket training, we reserve one core on each socket for the data loader thread, one core on each socket for an MPI communication thread, and rest of the cores on each socket for neural network computation. Please refer to the Methods section for more details on the implementation.

### Single-socket Experiments

In this experiment, we compare the oneDNN based implementation with our LIBXSMM based implementation running on multiple CPUs. We train the AtacWorks [2] neural network for 25 epochs on CLX, CPX, ROME, and ICX CPUs. We use a batch size of 64 with the code with oneDNN backend. With the LIBXSMM backend, batch size needs to be an integer multiple of available compute cores. Therefore, we use a batch size of 54 for CLX and CPX, a batch size of 63 for ROME, and a batch size of 78 for ICX. These batch sizes provide maximum training throughput for the corresponding CPUs.

Figure 3 shows the training time per epoch and achieved speedups on single-socket machines. The training time per epoch includes the evaluation time. We can observe that the LIBXSMM backend-based implementation on CLX achieves up to 6.86× speedup over the oneDNN library-based implementation on the same machine. Speedup increases on CPX as it has higher frequency and memory bandwidth compared to CLX. ICX is the next generation CPU with higher core count and provides further speedup as expected. Training time on ROME is slower than CLX, CPX, and ICX because ROME doesn’t support AVX-512 instructions. We also see significant speedup by training in BF16 precision without compromising the AUROC accuracy (Table 1). Overall, our implementation on CPX achieves 12.6× speedup compared to oneDNN based baseline implementation. This speedup is enabled not just by the use of modern CPUs with more cores and BF16 support, but especially due to our optimized implementation that uses the hardware well.

**Table 1.**
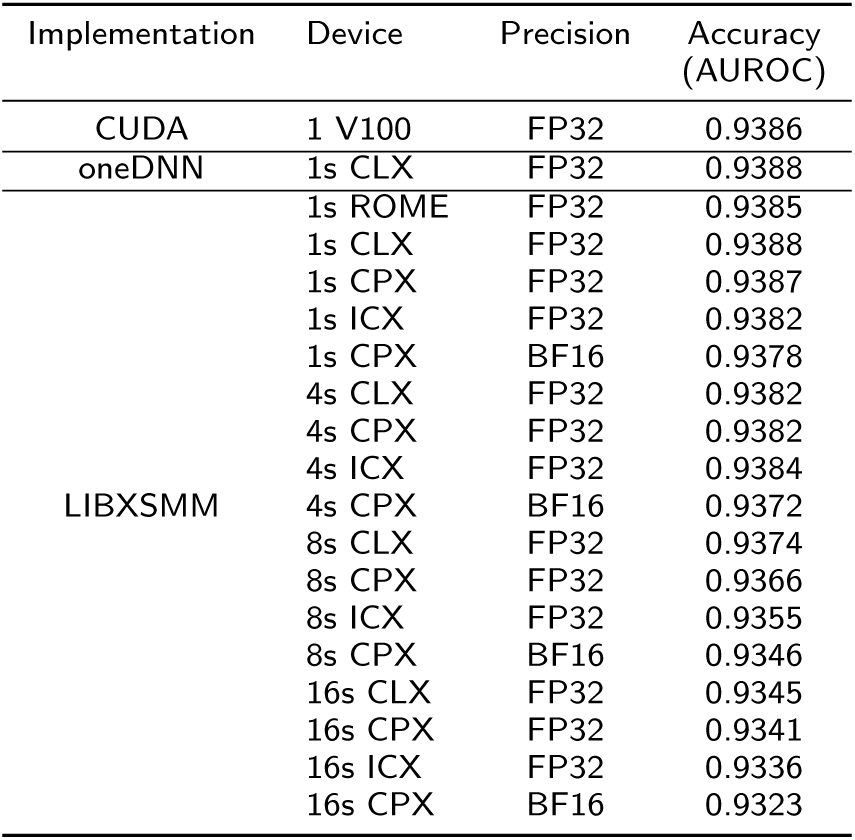
AtacWorks training accuracy results.

**Figure 3.**
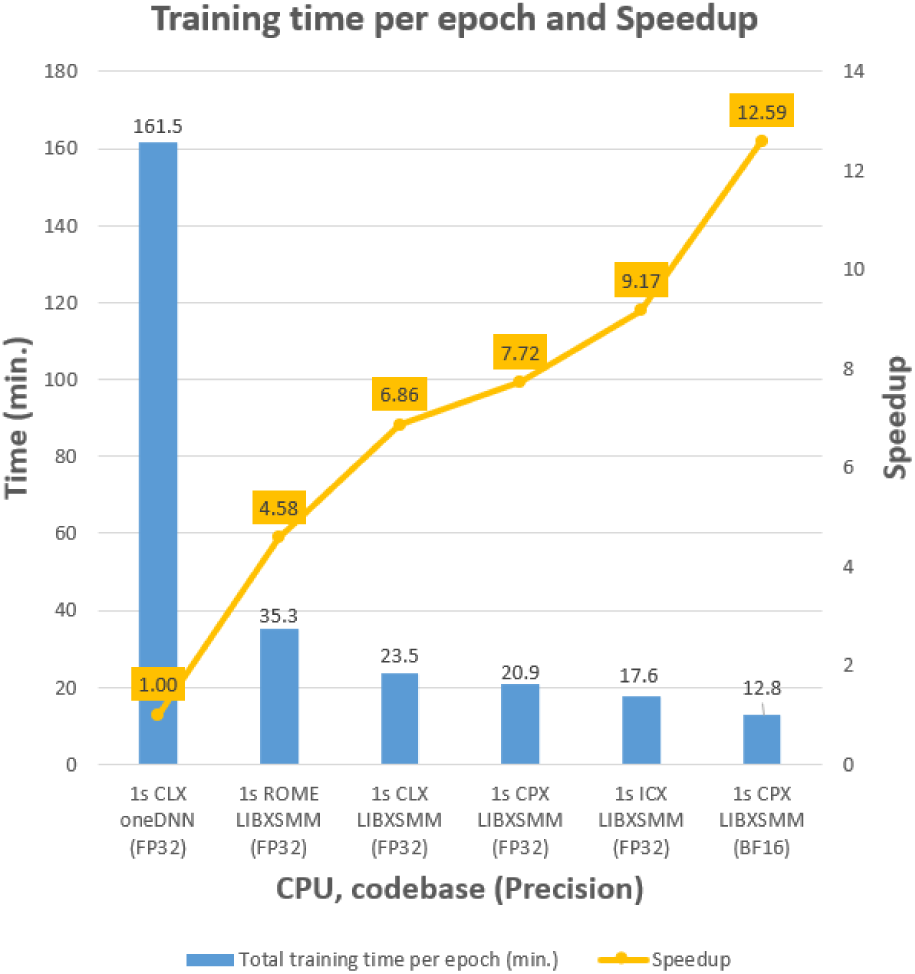
AtacWorks training time per epoch on one socket of CLX, CPX, ROME, and ICX.

### Multi-socket Experiments

In this set of experiments, we run the AtacWorks training experiment on multiple sockets of CPUs. By use of multiple sockets, we can significantly reduce the training time. It allows us to perform training on larger datasets. We also demonstrate the advantage of multi-socket CPU training in processing longer ATAC-seq signal track segments.

#### Multi-socket Training

We train the AtacWorks network for 25 epochs on multiple sockets of CLX, CPX, and ICX CPUs. For CLX and CPX experiments, we use 26 cores on each socket for computations and use a batch size that is an integer multiple of 26 and number of sockets. Therefore, we use a batch size of 104 on four sockets, 208 on eight sockets, and 416 on sixteen sockets. For ICX experiments, we use 38 cores on each socket for computations, and therefore, use a batch size of 152 on four sockets, 304 on eight sockets, and 608 on sixteen sockets. Figure 4 shows training time per epoch with multiple sockets of CLX, CPX, and ICX CPUs. We break down the total time into training and evaluation time in Figure 4 because the code that computes accuracy metrics such as mse, bce, AUROC, etc. in evaluation runs on a single-socket. The results show that our multi-socket implementation achieves nearly linear speedup for training time with the increase in the number of sockets.

**Figure 4.**
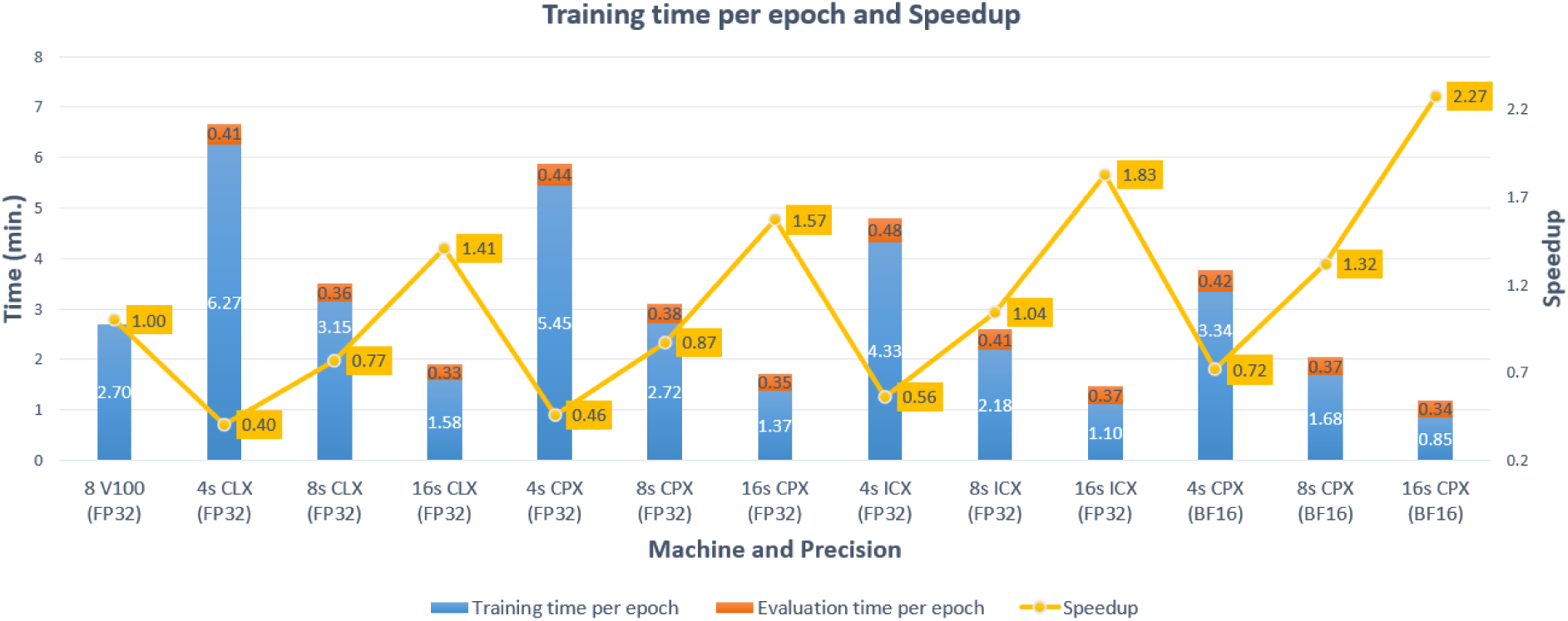
Training time per epoch and speedup with respect to 8 Nvidia V100s for AtacWorks training over 25 epochs on 8 Nvidia V100 and multiple sockets of CLX, CPX and ICX.

### Comparison with the state of the art

The state-of-the-art runtime as reported in the AtacWorks paper [2] is 2.7 minutes (162 seconds) per epoch for training on the DGX-1 box. A DGX-1 box [20] consists of 8 Nvidia V100 GPUs and a dual-socket host CPU. For a fair comparison, we compare the DGX-1 box with 16 sockets of CPU because they have a similar power envelop. Our optimized implementation running 16 sockets of CPX with BF16 consume only 1.19 minutes achieving a speedup of 2.27× over the DGX-1 box. Moreover, 16 sockets of CLX achieve 1.41× speedup, 16 sockets of CPX achieve 1.57× speedup, while 16 sockets of ICX with FP32 precision achieve 1.83× speedup over the DGX-1 box. In fact, even just 8 sockets of CPX with BF16 precision are 1.32× faster than the DGX-1 box. From Table 1, we can see that our implementation running on CPU-based systems achieves speedup over the original AtacWorks running on Nvidia V100 GPUs without any loss in accuracy.

#### Longer Signal Track Segment Experiment

The signal track size used so far allows us to find relationships spanning 60000 bases. Wider signal track segments may allow us to find relationships among bases that are farther apart. The maximum possible width of a signal track segment depends on the memory available on a machine. We can increase this width on CPUs due to large memory availability. Therefore, in order to demonstrate our implementation’s ability to apply deep learning on very long signal track segments, we also perform experiments on CPUs with input signal track segments of width 600, 000. We were not able to run this experiment on V100 GPUs, as the GPUs ran out of memory.

Due to the larger signal track segment size, there are fewer nonzero segments. As a result, we get 4191 signal track segments in the training set and 101 signal track segments in the validation set. Larger signal track segment size increases 1D dilated convolution layer sizes and activation tensor widths by 10×. We train AtacWorks on two sockets of CLX and used a batch size of 52. The training completed successfully in 407.2 minutes.

#### Multi-socket Experiment with Large Dataset

In order to evaluate whether our implementation performs well on a significantly larger dataset, we choose a larger real dataset with 293242 signal track segments of width 60000 (17.6 Billion values in total) in the training set instead of 32000 signal track segments of width 60000. We used 2400 monocytes cells signal and 2400 CD19 cells signal to make the clean high-coverage signal. The noisy set was composed of five different 50 cell subsamples of monocytes cells and five different 50 cell subsamples of CD19 cells. Thus, the training set in this experiment is approximately 9.16× (293242 vs. 32000) larger than the one used for all previous experiments. We also increase the validation set to 2520 signal track segments. Our per epoch training time is 14.5 minutes that is nearly 9.18× more than the training time on 16 sockets of CLX for 32000 signal track segments.Thus, 9.16× increase in dataset size resulted in nearly 9.18× increase in training time demonstrating linear scaling with the size of the dataset, as expected. Our total training and evaluation time for 25 epochs on 16 sockets of CLX is just 380 minutes. With this increased data size, we are also able to achieve a slightly higher training accuracy of 0.9390 in terms of the AUROC metric.

## Conclusion

We have accelerated the task of denoising and identifying chromatin accessibility via efficient implementation of deep learning on 1D data on multi-core CPUs. We present several methods of accelerating the AtacWorks toolkit, such as an optimized 1D dilated convolution layer with the LIBXSMM library’s TPPs, use of SIMD parallelism through AVX512 instructions, optimum thread placement, nearly linear scaling across multiple sockets and first ever BF16 based training without incurring any loss of accuracy. These optimizations accelerate the training by increasing code efficiency and achieve substantial speedups over previous implementations, thus, achieving the best throughput for AtacWorks.

Deep learning is increasingly finding applications in the field of genomics due to the increase in dataset sizes. We believe the extent of its application will depend on future availability and the cost of computing power. Efficient utilization of existing and future computing hardware will reduce the costs and allow researchers to conduct experiments at scale. 1D data is abundant in biology and deep learning is increasingly being used for processing it. Therefore, we believe that methods presented in this work, such as optimized 1D convolution layer, ability to perform deep learning on long 1D sequences, and BF16 computations, can be applied to many other problems in biology. In addition, our generic implementation of optimized 1D convolution layer can readily be used in other applications.

## Methods

In this section, we briefly describe the techniques used to accelerate the AtacWorks toolkit using modern multicore CPUs. First, we demonstrate that the current method for performing 1D convolution on CPUs is inefficient and describe our optimized convolution layer that utilizes the CPUs efficiently. Second, we describe how we optimize scaling to multiple threads on a single CPU socket and to multiple CPU sockets while overlapping data loading with computation. These optimizations allow efficient AtacWorks training on multiple sockets of multi-core CPUs allowing us to process large datasets. Finally, we describe how we use reduced precision (BF16) to get further speedup.

### Optimized Convolution Layer

Most convolution layers in AtacWorks have 15 channels, 15 filters, a filter width of 51, and a dilation parameter of 8. These layers also have 1D input and output of widths 60400 and 60000, respectively. For these parameters, current implementations for 1D dilated convolution layer are inefficient and significantly underutilize CPU resources. We demonstrate this by creating a single convolution layer in the PyTorch framework and timing its forward pass and backward pass for 20 iterations. Specifically, we time *Y = net.forward(X)* method for the forward pass, and *Y.sum().backward()* method for the backward pass. This method of measuring the execution time may have some framework overhead compared to measuring execution time during a training run. Still, it provides a good yardstick for performance comparisons and efficiency assessment of various implementations. For these experiments, we use one OpenMP thread per core and perform multi-threading by distributing across the threads the signal track segments that are within a batch. We can observe from Supplementary Table S2 that the PyTorch layer with default backend can only achieve less than 1% of peak machine performance. If we replace the default PyTorch backend with oneDNN based backend for convolutions, it can achieve 19.9% and 4.1% of peak machine performance for forward pass and backward pass, respectively. While this is better utilization than PyTorch, it is still far from the peak efficiency. With oneDNN backend, PyTorch convolution layers take more than 90% of the total AtacWorks training time. Thus, further improvements are necessary to increase the efficiency of the 1D convolution layer.

Here, we briefly describe our accelerated 1D dilated convolution layer. We provide a more detailed description in Supplementary Note 1. To create our accelerated 1D dilated convolution layer, we create an efficient and generic C++ implementation of three compute kernels – the forward pass, the backward-by-data pass, and the backward-by-weight pass – and integrate them into the PyTorch framework by making a PyTorch C++ extension for the convolution layer. To implement an efficient and generic version of the forward and the backward pass kernels, we use the LIBXSMM library’s copy, transpose, GEMM, and BRGEMM TPPs [16]. TPPs leverage Just-in-time (JIT) code generation technology to generate platform-specific code. Therefore, it can generate AVX-512 instructions at FP32 precision for CLX CPUs. Using LIBXSMM’s TPPs, we create a cache blocked implementation with AVX-512 (FP32) instructions to efficiently utilize CLX CPU’s resources. The third column in Supplementary Table S2 shows the flops, efficiency, and speedup results of the PyTorch layer that uses our efficient implementation as backend. This optimized code achieves up to 74.3% of machine peak flops in the forward pass and up to 55.7% of the machine peak flops in the backward pass. Additionally, it is nearly 107× and 86× faster in the forward pass and backward pass, respectively, over the PyTorch baseline layer with unoptimized default backend.

### Multi-threading and Multi-socket optimizations

For a stand alone 1D convolution layer, using 1 thread per core is efficient. However, this strategy is not efficient during end-to-end training of a neural network where in every epoch large amounts of data are loaded from disk to memory. If we do not reserve some cores for data loading then compute cores may idle during data loading. Therefore, we use one core exclusively for loading data and use the remaining cores for computations. We read data using the PyTorch framework’s *DataLoader()* workers that are independent processes. Supplementary Figure S3 shows this configuration, and it allows efficient utilization of resources by reducing conflicts between *DataLoader()* worker threads and OpenMP compute threads. Additionally, we also keep the batch size as an integer multiple of the number of OpenMP threads so that we can distribute equal number of signal track segments to all the threads.

For multi-socket training, we perform communication across sockets using MPI [23] through oneCCL library [22]. On each socket, we reserve one core for *DataLoader()*, one core for communication across sockets using MPI, and 26 cores for OpenMP threads for computation. Supplementary Figure S4 shows the thread placement when using two sockets of 28 core CPUs.

### BFloat16 Optimization

Our third set of optimizations focus on the use of reduced precision training of neural networks. Some processors like Intel’s Cooper Lake (CPX) support AVX-512 BF16 instructions. With half the number of bits used compared to FP32, use of BF16 can theoretically double the performance without significant accuracy loss in deep learning applications [24, 25]. To leverage BF16 compute, we first write a BF16 version of the 1D dilated convolution layer. We can easily convert the TPP based FP32 precision source code into a BF16 precision source code. However, there are some constraints. For example, our BF16 source code requires an even number of filters, channels, and elements along the input tensor width. Therefore, we use 16 filters and 16 channels in our BF16-based convolution layer compared to 15 filters and 15 channels used for FP32 precision. Our BF16-based implementation achieves 64% and 44.6% efficiency for forward pass and backward pass, respectively, compared to the peak CPX performance when using BF16 instructions (Supplementary Table S2). This results in 199× and 148.3× speedups for forward pass and backwards pass, respectively, over the baseline FP32-based PyTorch layer running on CLX.

A BF16 precision convolution layer alone is not enough to perform end-to-end training in BF16 precision. We need the entire network to perform computations in BF16 precision. If some layers have BF16 compute while others have FP32, then there can be multiple tensor conversion operations in the computational path. Data conversion operations are time-consuming, and they reduce the benefits of BF16 compute. Therefore, we must ensure that, during training, the majority of the computation is in BF16 precision. To achieve this, we make a BF16 precision-based ReLU layer. We also convert the input tensor into BF16 precision and keep almost all operations in BF16. The loss layers are the only layers that don’t work on BF16 precision and data must be converted to FP32 for them. Supplementary Figure S5 shows an updated Resblock with all tensors and layers in BF16.

## Supporting information

Supplemental section

## Availability of data and materials

AtacWorks CPU code is available as a patch to original AtacWorks code. The patch location and instructions are available at https://github.com/IntelLabs/Trans-Omics-Acceleration-Library/tree/ATAC-Seq/applications/ATAC-Seq. Follow the instructions in *patch instructions.md* file to make a training run.

## Ethics approval and consent to participate

Not applicable.

## Competing interests

NC, SM, DK, AH, EG, BZ, MA and BK are employees of Intel Corporation.

## Authors’ contributions

NC led the software implementation under the guidance of SM and DK. AK, EG, BZ and MA created the LIBXSMM TPPs. All authors contributed to design of algorithm, experiments and manuscript preparation. All authors read and approved the final manuscript.

## Optimization Notice

Software and workloads used in performance tests may have been optimized for performance only on Intel microprocessors. Performance tests, such as SYSmark and MobileMark, are measured using specific computer systems, components, software, operations and functions. Any change to any of those factors may cause the results to vary. You should consult other information and performance tests to assist you in fully evaluating your contemplated purchases, including the performance of that product when combined with other products. For more information go to http://www.intel.com/performance. Intel, Xeon, and Intel Xeon Phi are trademarks of Intel Corporation in the U.S. and/or other countries.

## Notes

### Competing Interest Statement

All authors are employees of Intel Corporation.

