## Supplemental section for "Accelerating Identification of Chromatin Accessibility from noisy ATAC-seq Data using Modern CPUs"

### Supplementary Data

**Table S1** Architectural specifications of four processors: Cascade Lake, Cooper Lake, Ice Lake, and ROME which were used for the experiments.

| | Intel® Xeon® Platinum 8280 Cascade Lake | Intel® Xeon® Platinum 8380H Cooper Lake | Intel® Xeon® Platinum 8380 Ice Lake | AMD® EPYC® 7742 ROME |
|---|---|---|---|---|
| Cores / Socket | 28 | 28 | 40 | 64 |
| Threads / Cores | 2 | 2 | 2 | 2 |
| AVX register width (bits) | 512, 256, 128 | 512, 256, 128 | 512, 256, 128 | 256, 128 |
| Vector Processing Units (VPU) | 2/Core | 2/Core | 2/Core | 2/Core |
| Base Clock Frequency (GHz) | 2.7 | 2.9 | 2.4 | 2.25 |
| L1D, L2 Cache (KB) | 32, 1024 | 32, 1024 | 48, 1280 | 32, 512 |
| L3 Cache (MB) / Socket | 38.5 | 38.5 | 60 | 256 |
| DRAM (GB) / Socket | 96 | 192 | 128 | 128 |
| Bandwidth (GB/s) / Socket | 128 | 154 | 205 | 205 |
| Compiler Version | GCC v. 8.3.0 | GCC v. 8.3.0 | GCC v. 8.3.0 | GCC v. 8.3.0 |

#### Supplementary Note1: Design of Our Efficient 1D Dilation Convolution Layer

We implement the forward pass and the backward-by-data pass of the 1D dilated convolution layer using BRGEMM kernel of the LIBXSMM library. The backward-by-weight pass kernel is implemented using small GEMM kernels. We do not implement the bias calculation of the forward and the backward pass but instead use the PyTorch framework's implementation. BRGEMM kernel multiplies two matrix blocks $A_i \in \mathbb{R}^{m\times k}$ and $B_i \in \mathbb{R}^{k\times n}$ and reduces the partial results to a block $C_j \in \mathbb{R}^{m\times n}$ of a tensor C. The blocks $A_i$ and $B_i$ can be taken from any position in the larger A and B input tensors. BRGEMM kernel needs the following arguments: $(i)$ Arrays of pointers for the $A_i$ and $B_i$ blocks to be multiplied, $(ii)$ a pointer to the output block $C_j$, $(iii)$ Size of blocks, $(iv)$ the number $(l_{br})$ of the blocks to be multiplied and $(v)$ the scaling parameters $\alpha$ and $\beta$. Equation (3) shows the batch reduce GEMM operation.

$$C_j = \beta * C_j + \alpha * \sum_{0}^{l_{br}-1} A_i * B_i \tag{1}$$

Figure S1 illustrates batch-reduce GEMM operation with two-dimensional tensors. As shown in the figure, we can choose matrix blocks from any place in the tensor by specifying pointers and block sizes. It is also possible for the matrix blocks to overlap. In the following subsections, we present the forward pass, the backward-by-data pass, and the backward-by-weight pass algorithms in more detail.

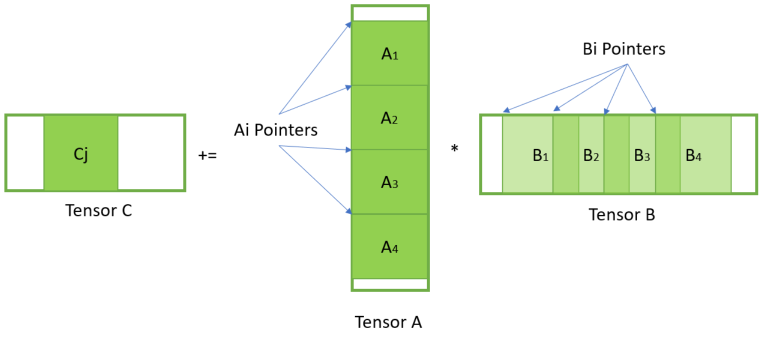


**Figure S1** Example of Batch-reduce GEMM with 2D tensors. Blocks at $A_i$ and $B_i$ positions get multiplied and reduced to the $C_j$ block.

##### 0.1 Forward Pass

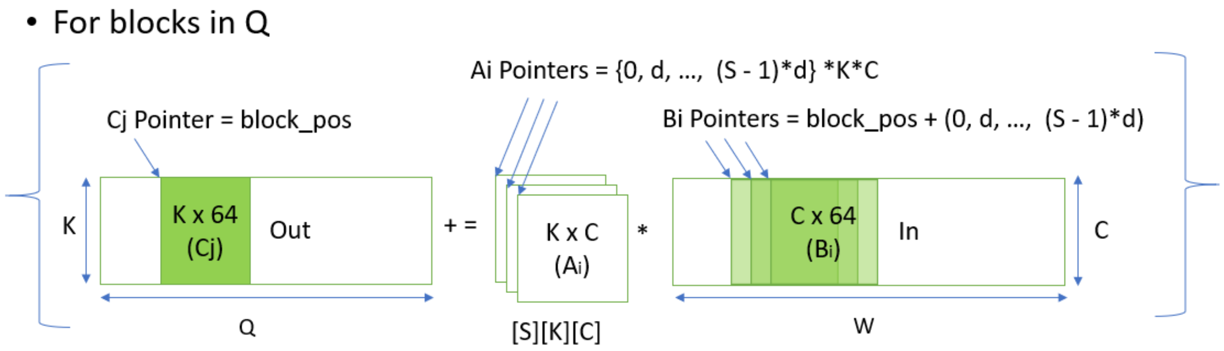


**Figure S2** Forward pass kernel using batch-reduce GEMM. We multiply $A_i$ blocks from the weight tensor and $B_i$ blocks from the input tensor. The result is reduced into $C_j$ block in the output tensor. Cache blocking occurs along the width dimension.

To implement the forward pass, we first make some changes in the weight tensor layout to convert the forward pass computation into a matrix multiplication. We change the layout of the weight tensor from $(K, C, S)$

to $(S, K, C)$. Consequently, the 1D dilated convolution can be described by a series of $S$ GEMM operations explained in Algorithm 1.

**Algorithm 1** Forward pass using GEMM operations

**Inputs:** $In \in \mathbb{R}^{C \times W}$, $Weight \in \mathbb{R}^{S \times K \times C}$, $d \in \mathbb{R}$
**Output:** $Out \in \mathbb{R}^{K \times Q}$

**procedure** FORWARD PASS$(Out, In, Weight, d)$
  **for** $s = 0, 1, ..., S-1$ **do**
    $Out[:, :] \mathrel{+}= \mathbf{GEMM}(Weight[s, :, :], In[:, (d*s) : (d*s+Q)])$
  **return** $Out$ ▷ The output tensor

Once we can express the algorithm in terms of GEMM operations, we can convert it into BRGEMM operations. We can replace the for loop of Algorithm 1 with a BRGEMM computation. A matrix problem-size suitable for the LIBXSMM library with matrix dimensions $m, n, k$ is that satisfies $(mnk)^{1/3} <= 64$. For such sizes, the LIBXSMM library automatically employs an efficient utilization of the cache hierarchy. Thus, we employ blocking along the input width dimension and perform BRGEMM operation on the blocks. In all our kernels, we keep the block length equal to 64 elements along the width dimension. Block length of 64 ensures that one dimension of the GEMM problem size remains within the LIBXSMM library's constraints. In our implementations, the other two dimensions are defined by the number of channels $C$ and the number of filters $K$ parameters. Thus, we achieve a highly cache efficient implementation whenever $(C*K)^{1/2} <= 64$. Additionally, the LIBXSMM library's GEMM kernels maintain good efficiency as long as the value of $(C*K)^{1/2}$ is not drastically higher than 64. Algorithm 2 and figure S2 show the forward pass computation using BRGEMM kernel.

**Algorithm 2** Forward pass using BRGEMM operation

**Inputs:** $In \in \mathbb{R}^{C \times W}$, $Weight \in \mathbb{R}^{S \times K \times C}$, $d \in \mathbb{R}$
**Output:** $Out \in \mathbb{R}^{K \times Q}$

**procedure** FORWARD PASS$(Out, In, Weight, d)$
  **for** $pos = 0 \ to \ Q \ in \ steps \ of \ 64$ **do** ▷ Cache blocking
    **for** $s = 0, 1, ..., S-1$ **do** ▷ Generate pointers
      $A_{ptrs}[s] = Pointer \ to \ Weight[s, 0, 0]$
      $B_{ptrs}[s] = Pointer \ to \ In[0, (pos + s*d)]$
    $\mathbf{BRGEMM}(A_{ptrs}, B_{ptrs}, Pointer \ to \ Out[0, pos], S)$
  **return** $Out$ ▷ The output tensor

#### 0.2 Backward-by-Data Pass

In the backward-by-data pass, we first change the weight tensor layout from $(K, C, S)$ to $(S, C, K)$. Similar to the forward pass, data gradient $(Grad_d \in \mathbb{R}^{C \times W})$ computation can be converted into a matrix multiplication of weights and output gradient $(Grad_{out} \in \mathbb{R}^{K \times Q})$. The backward data pass algorithm is very similar to the forward pass. We again employ the cache blocking along the width dimension with a block size of 64. We zero pad the gradient output $(Grad_{out})$ wherever needed. Algorithm 3 implements the backward data kernel using BRGEMM operation.

**Algorithm 3** Backward-by-data pass using BRGEMM operation

**Inputs:** $Grad_{out} \in \mathbb{R}^{K \times Q}$, $Weight \in \mathbb{R}^{S \times C \times K}$, $d \in \mathbb{R}$
**Output:** $Grad_d \in \mathbb{R}^{C \times W}$

**procedure** BACKWARD-BY-DATA PASS$(Grad_d, Grad_{out}, Weight, d)$
  **for** $pos = 0 \ to \ W \ in \ steps \ of \ 64$ **do** ▷ Cache blocking
    **for** $s = 0, 1, ..., S-1$ **do** ▷ Generate pointers
      $A_{ptrs}[s] = Pointer \ to \ Weight[s, 0, 0]$
      $B_{ptrs}[s] = Pointer \ to \ Grad_{out}[0, pos - (S-1-s)*d]$
    $\mathbf{BRGEMM}(A_{ptrs}, B_{ptrs}, Pointer \ to \ Grad_d[0, pos], S)$
  **return** $Grad_d$ ▷ Data gradient

#### 0.3 Backward-by-Weight Pass

We utilize small GEMM operations in the backward-by-weight pass implementation. We again do cache blocking along the width dimension with a block size of 64. The backward-by-weight pass module is less efficient than the other modules because the data blocks cannot be kept in the cache for a long time. Additionally, the weight tensor must be shared across multiple threads when multithreading on the batch dimension ($N$). Algorithm 4 implements the backward weight pass using small GEMM operations.

**Algorithm 4** Backward weight pass using small GEMM operations

**Inputs:** $Grad_{out} \in \mathbb{R}^{K \times Q}$, $In \in \mathbb{R}^{C \times W}$, $d \in \mathbb{R}$
**Output:** $Grad_w \in \mathbb{R}^{S \times C \times K}$

```
1: procedure BACKWARD WEIGHT PASS(Grad_w, Grad_out, In, d)
2:     for pos = 0 to Q in steps of 64 do                                  ▷ Cache blocking
3:         for s = 0, 1, ..., S − 1 do
4:             Grad_w[s, :, :] += GEMM(In[:, (pos + s ∗ d) : (pos + s ∗ d + 64)], transpose(Grad_out[:, pos : (pos + 64)]))
5:     return Grad_w                                                       ▷ Weight gradient
```

**Table S2** The efficiency of 1D dilated convolution layer's forward pass and backward pass. FP32 results are obtained on a $28$ core single-socket Cascade Lake CPU. We also used a $28$ core Cooper Lake CPU to get BF16 results. Peak machine performance of Cascade Lake is $4.3$ TeraFLOPS (FP32), Cooper Lake is $9.32$ TeraFLOPS (BF16). We time a single convolution layer network and take an average of 20 iterations. For these experiments, we use one OpenMP thread per core and perform multi-threading along the batch dimension of the input tensor. Therefore, we use $28$ OpenMP threads for computation on a $28$ core single-socket CPU.

| Compute kernel | Without oneDNN (CLX) GFlops — Effi. — Speedup | oneDNN Backend (CLX) GFlops — Effi. — Speedup | LIBXSMM Backend (CLX) GFlops — Effi. — Speedup | LIBXSMM Backend (BF16) (CPX) GFlops — Effi. — Speedup |
|---|---|---|---|---|
| Forward pass | 30 — 0.7% — 1.0x | 857 — 19.9% — 28.6x | 3197 — 74.3% — 106.6x | 5969 — 64% — 199.0x |
| Backward pass | 28 — 0.7% — 1.0x | 178 — 4.1% — 6.4x | 2396 — 55.7% — 85.6x | 4153 — 44.6% — 148.3x |

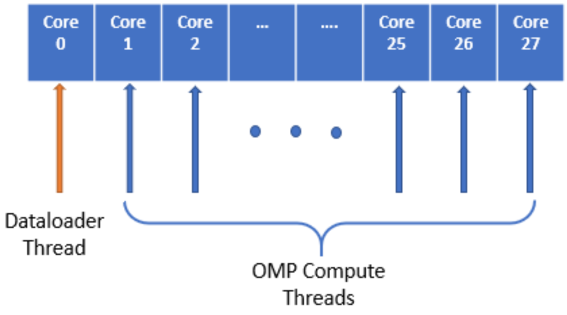


**Figure S3** Dataloader worker and OMP compute thread positioning on a single-socket 28 core CPU.

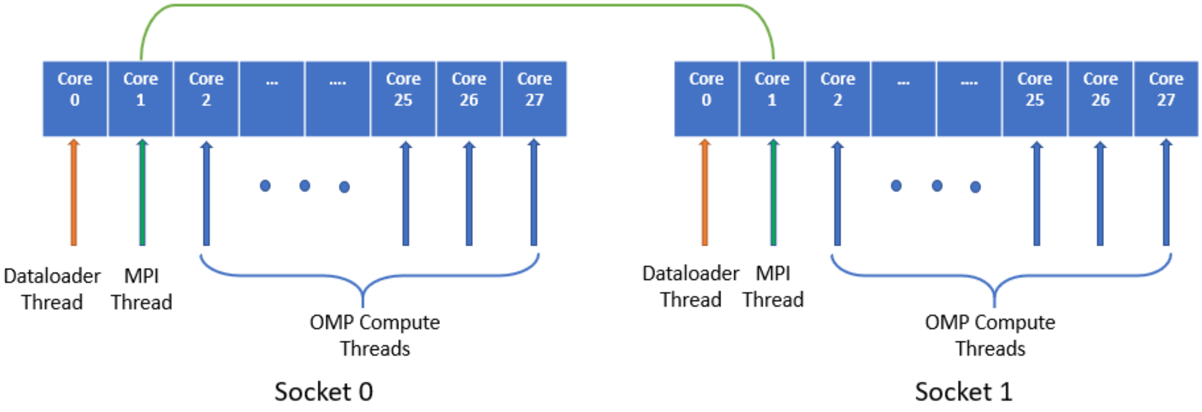


**Figure S4** Dataloader, MPI and OMP compute thread positioning for a multi-socket run.

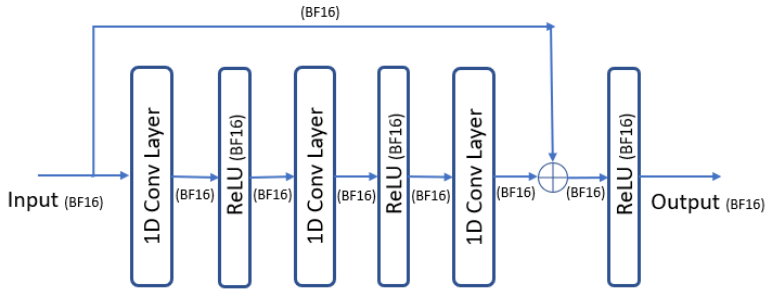


**Figure S5** Residual block in BFloat16 precision.
